# MXene–Protein Corona Interfaces for Molecular Profiling of Alzheimer’s Disease

**DOI:** 10.64898/2026.05.14.725150

**Authors:** Samantha Velazquez, Matthew C. Juber, Dalton Brindley, Anupma Thakur, Babak Anasoori, Edward Lau, Ali Akbar Ashkarran

**Affiliations:** Department of Biology, University of Colorado Colorado Springs, Colorado Springs, CO 80918, USA; Department of Medicine, University of Colorado School of Medicine, Aurora, CO 80045, USA; Department of Physics and Energy Science, University of Colorado Colorado Springs, Colorado Springs, CO 80918, USA; School of Materials Engineering, Purdue University, West Lafayette, IN 47907, USA; School of Mechanical Engineering, Purdue University, West Lafayette, IN 47907, USA; BioFrontiers Center, University of Colorado Colorado Springs, Colorado Springs, CO 80918, USA

**Keywords:** Nanomaterials’ protein corona, Alzheimer disease, Ti_3_C_2_T*_x_* MXene, Proteomics, Biomarker discovery

## Abstract

The protein corona (PC) that forms on the surface of nanomaterials upon contact with biological fluids provides a molecular snapshot of the host’s physiological and pathological state. Here, we investigate two-dimensional (2D) titanium carbide (Ti_3_C_2_T*_x_*) MXene nanosheets as nanobiointerfaces for capturing Alzheimer’s disease (AD)–associated plasma protein signatures. Ti_3_C_2_T*_x_* MXene flakes were incubated with plasma from clinically diagnosed AD patients and age-matched healthy controls (HC), leading to the formation of Ti_3_C_2_T*_x_* MXene–PC complexes. Physicochemical characterization using dynamic light scattering, zeta potential analysis, and transmission electron microscopy revealed disease-dependent changes in hydrodynamic size, surface charge, and PC profile. Proteomic analysis of the isolated PC layers quantified 1,611 proteins without prior fractionation, demonstrating effective enrichment of low-abundance plasma components. Principal component analysis (PCA) revealed consistent separation between AD- and HC-derived Ti_3_C_2_T*_x_* MXene–PC proteomes despite inter-individual heterogeneity. Differential abundance analysis identified selective enrichment of heterogeneous nuclear ribonucleoproteins (hnRNPs), annexins, and inflammatory mediators in AD-derived PC, implicating dysregulated RNA metabolism, membrane stress responses, and immune activation, hallmark processes in AD pathology. Our findings demonstrate that Ti_3_C_2_T*_x_* MXene–PC interfaces act as selective molecular filters that reshape the detectable plasma proteome, enabling disease-associated molecular phenotyping and establishing a versatile nanointerface-driven framework for uncovering AD-related plasma signatures, providing a foundation for future translational diagnostic development.

## Introduction

Alzheimer’s disease (AD) is a progressive neurodegenerative disorder characterized by cognitive decline, neuronal loss, and the pathological accumulation of amyloid-β (Aβ) plaques and hyperphosphorylated tau (p-tau) tangles in the brain.^1-3^ Despite decades of research, early and accurate diagnosis of AD remains a major clinical challenge. Conventional diagnostic approaches, including neuroimaging, cerebrospinal fluid (CSF) analysis, and neuropsychological testing, are either invasive, costly, or lack the sensitivity required for detecting early-stage disease.^4-8^ Blood-based biomarker detection has emerged as a promising alternative. However, the extremely low abundance of AD-associated proteins in plasma and the high background of nonspecific proteins hinder reliable quantification. For example, phosphorylated tau (p-tau) proteins, key indicators for AD, are typically present at very low levels (<1 pg/mL) in the bloodstream, which makes it difficult to detect them accurately, especially with traditional diagnostic methods.^9^ This is mainly because of the wide dynamic range of the plasma proteome in protein abundance (i.e., from g/L to pg/L) which overshadows and obscures the detection and quantification of lower-abundance proteins relevant to AD diagnosis due to the presence of other highly abundant proteins (e.g., albumin, immunoglobulin, and transferrin) in human plasma.^10-11^ Therefore, developing techniques that can effectively capture and analyze low-abundance proteins with high AD diagnostic potential still remains highly challenging.

The protein corona (PC) is a layer of biomolecules (mainly proteins) that forms around the surface of nanomaterials after contact with a biomolecular source such as serum or plasma.^12-14^ The PC layer initially consists of proteins with higher abundancy and lower affinities (e.g., albumin) in the biological fluid; however, in a dynamic process, the initial PC layer will be replaced over time with higher affinity and lower abundant proteins (known as hard PC), which strongly adsorb on the surface of the NPs and remain stable for longer periods of time. ^15-17^ In this regard, NPs’ PC has the potential to enrich the human plasma proteome due to the dynamic nature of the PC formation on the surface of the nanomaterials.^18^ One of the key aspects of PC in biomarker discovery is enrichment of low-abundance biomarkers, which are present in human plasma at very low concentrations and can become enriched in the PC layer.^19-20^ This enrichment facilitates the detection and identification of rare biomarkers that might be challenging to detect directly from the complex biological matrix (i.e., human plasma).^21^

Recent advances in nanomaterials have opened new pathways for sensitive biomolecular detection through modulation of the nano–bio interface. The specific proteins that adsorb onto the surface of nanomaterials depend on various factors, including the size, shape, surface charge, and surface chemistry of the nanomaterials, as well as the composition of the surrounding biological fluid.^22^ Two-dimensional (2D) nanomaterials have attracted substantial attention in biosensing and diagnostic applications due to their high surface area, tunable electronic properties, and strong biomolecular interactions.^23-27^ In particular, graphene and its derivatives have been widely explored for AD detection, including electrochemical and optical sensing of biomarkers such as amyloid-β and tau proteins.^28-34^ These platforms have demonstrated promising sensitivity; however, many rely on targeted detection strategies and surface functionalization, which may limit their ability to capture the broader complexity of disease-associated proteomic changes.^35-40^

In this context, MXenes, a large class of 2D transition metal carbides, nitrides, and carbonitrides, offer unique advantages including intrinsically hydrophilic surfaces, rich surface terminations (e.g., –OH, –O, –F), and strong biomolecule adsorption capacity, which have led to increasing interest in their use as biointerfaces for protein interaction and adsorption studies. These properties make MXenes particularly promising candidates for PC–based, untargeted molecular profiling approaches.^41-46^ For instance, the increased surface reactivity and adsorption efficiency enable more effective biomarker sequestration, particularly under physiological conditions where target proteins exist at sub-nanomolar concentrations.^47^ These unique properties make MXenes particularly suitable for constructing biosensing interfaces capable of efficient biomolecular interaction and capture disease disease-related biomarkers.^48-49^ Importantly, the large specific surface area and planar morphology of 2D MXenes act synergistically with the dynamic nature of PC formation. This synergistic interplay between the 2D MXene surface and PC dynamics enables deeper sampling of the plasma proteome, enhancing the capture of disease-specific proteins (e.g., p-tau and amyloid-β).^50^ Such properties position MXenes as a powerful platform for ultrasensitive detection and precision nanodiagnostics, particularly for capturing subtle molecular signatures associated with AD. However, despite the growing interest in MXene-based biointerfaces, their use in PC–mediated disease profiling remains largely unexplored.

In this work, we use Ti_3_C_2_T*_x_* MXene as the nanomaterial platform due to its unique combination of physicochemical and biological properties that make it particularly suitable for plasma proteomic applications including^44-45^ (i) high electrical conductivity and surface reactivity of MXenes which facilitate rapid charge redistribution and strong electrostatic interactions with biomolecules, enabling stable and reproducible PC formation^51-52^, (ii) hydrophilic, surface-terminated structure which allows controllable surface chemistry and functionalization, ensuring compatibility with aqueous biological media and minimizing nonspecific aggregation^53-55^, and (iii) MXenes’ high biocompatibility and colloidal stability in physiological environments, as demonstrated in several recent in vitro and in vivo studies, which makes MXenes clinically relevant nanomaterials for biosensing and diagnostic applications.^48, 56-57^ An overview of the experimental workflow is illustrated in **Figure 1**. Plasma samples from AD patients and healthy controls were incubated with Ti_3_C_2_T*_x_* MXene nanosheets to form PC complexes, followed by washing and isolation of MXene–PC structures. Upon incubation with plasma from AD patients and healthy controls, MXene nanosheets spontaneously acquire a biomolecular coating that reflects the composition of the surrounding proteome. These Ti_3_C_2_T*_x_* MXene–PC complexes are subsequently subjected to comprehensive physicochemical characterization and label-free quantitative proteomic analysis to elucidate group-specific molecular fingerprints. This integrative approach offers a versatile framework for nanointerface-driven molecular profiling in AD, demonstrating the potential of Ti_3_C_2_T*_x_* MXene–PC systems as minimally invasive platforms for decoding disease-specific plasma alterations and precision neurodiagnostics.

**Figure 1:**
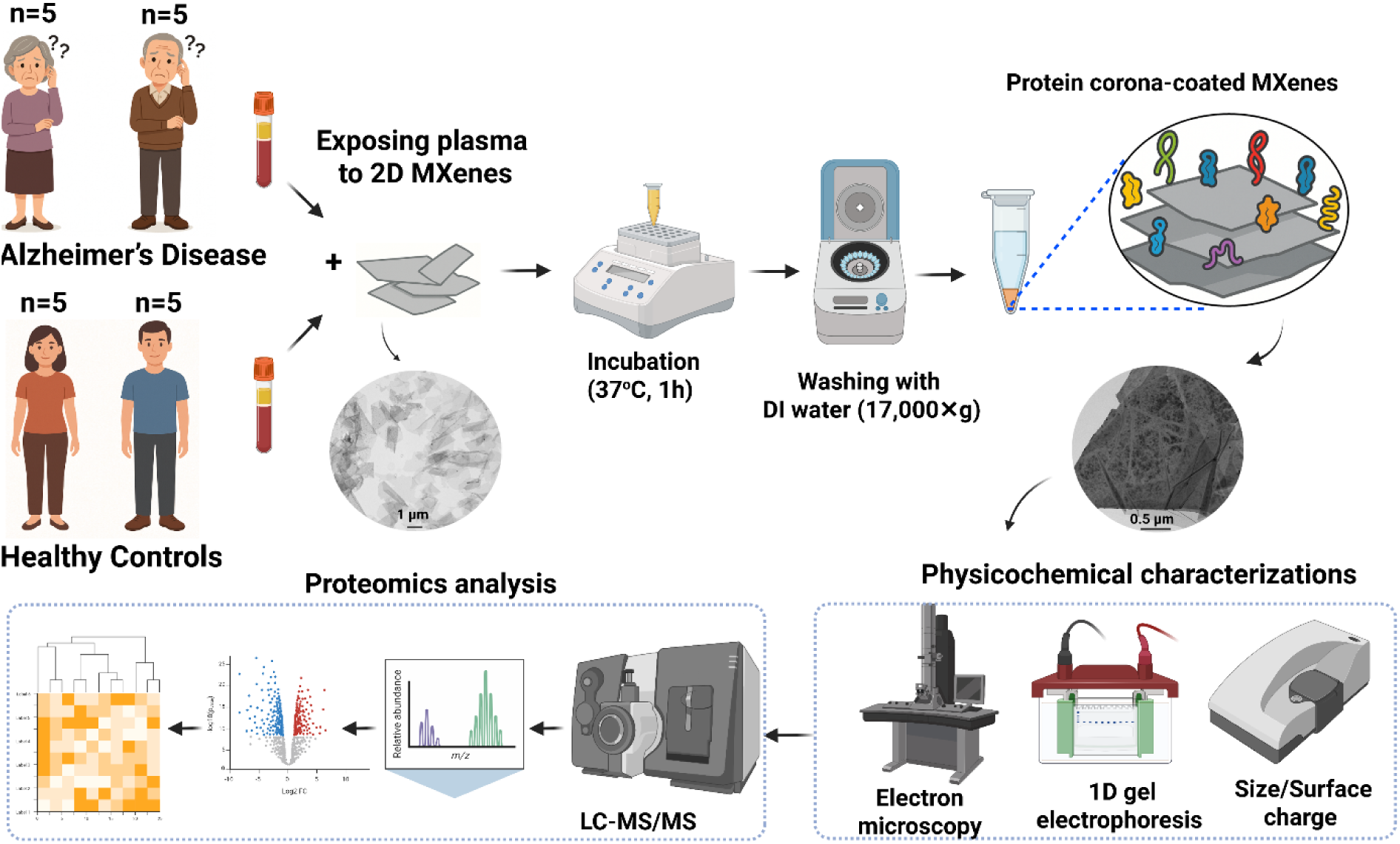
Overall workflow and experimental design for MXene-based AD detection using plasma PC profiling.

## Experimental details

### Materials

All reagents were used as received without further purification. Lithium chloride (LiCl, ≥98%), 28.4 M hydrofluoric acid (HF), and hydrochloric acid (HCl, 37%) were purchased from Thermo Fisher. Ethanol (≥99.5%) and deionized water (18.2 MΩ·cm) were used throughout the synthesis and washing steps. Ammonium bicarbonate, urea, dithiothreitol, iodoacetamide, and acetonitrile were purchased from Sigma-Aldrich. Trypsin was obtained from Promega, and formic acid was purchased from Thermo Scientific. AD and healthy human plasma samples ordered from Precision for Medicine and were diluted to a final concentration of 55% using DI water.

### Synthesis of Ti_3_C_2_T_*x*_ MXene

To synthesize Ti_3_C_2_T*_x_* MXene, 1 g of Ti_3_AlC_2_ MAX synthesized with excess metals (Ti and Al) was first washed using 9 M HCl for 18 h at room temperature to remove intermetallic impurities and mixed with an etchant solution (6:3:1 mixture (by volume) of 12 M HCl, deionized water, and 28.4 M HF before stirring at 400 RPM for 24 h at 35 °C.^58^ The etched Ti_3_C_2_T*_x_* MXene was then washed with deionized water via repeated centrifugation at 3234 RCF (4-5 cycles with ∼200 mL of deionized water) until the supernatant reached pH ∼6. Furthermore, for delamination, the etched multilayered Ti_3_C_2_T*_x_* MXene sediment was then added to the LiCl solution, typically 50 mL per gram of the starting etched powder. The mixture of LiCl and multilayer MXene was then stirred at 400 RPM for 1 h at 65 °C under constant argon gas flow. The mixture was then washed with deionized water via centrifugation at 3234 RCF for 5, 10, 15, and 20 minutes. Then, the final mixture was vortexed for 30 minutes, followed by centrifugation at 2380 RCF for 30 minutes to ensure the Ti_3_C_2_T*_x_* MXene solutions were single-to-few-layered flakes.

### PC formation on the surface of the MXenes

For PC formation, 2D Ti_3_C_2_T*_x_* MXenes were incubated with diluted (55%) plasma samples from healthy controls and AD patients so that the MXenes’ and human plasma final concentrations were 0.1 mg/ml and 55% in a 1 ml batch, respectively. The solution was then mixed, vortexed well, and incubated for 1h at 37 °C at a constant 1000 rpm agitation. To remove unbound and plasma proteins only loosely attached to the surface of MXenes, protein-MXene complexes were then centrifuged at 17,000× g for 20 minutes, the collected MXenes’ pellets were washed twice more with DI water under the same conditions, and the final pellet was collected for preparation for further analysis.

### LC-MS/MS sample preparation

The Ti_3_C_2_T*_x_* MXene-PC pellets obtained from various healthy and AD plasma samples were subjected to a standardized proteomic digestion protocol optimized for mass spectrometry based on our previous findings.^59^ Each PC-coated Ti_3_C_2_T*_x_* MXenes’ pellet was resuspended in 25 µL of buffer containing 50 mM ammonium bicarbonate and 1.6 M urea to facilitate protein solubilization and denaturation. Reduction was achieved by adding 2.5 µL of dithiothreitol (DTT, Pierce BondBreaker, neutral pH) to a final concentration of 10 mM. Samples were incubated at 37 °C for 60 minutes using a Thermomixer. Following incubation, samples were brought to room temperature, vortexed briefly, and centrifuged to collect the contents. Alkylation was performed by adding 2.7 µL of freshly prepared iodoacetamide (IAA) to a final concentration of 50 mM, followed by incubation in the dark at room temperature for 1 hour.

To quench excess IAA, 3 µL of DTT (10 mM final concentration) was added, followed by a 15-minute incubation at room temperature. Subsequently, sequencing-grade trypsin (Promega), diluted in protein digestion buffer, was added at a 1:50 trypsin-to-protein ratio. Enzymatic digestion proceeded overnight (∼16 h) at 37 °C in a Thermomixer. The next day, the digested samples were cooled to room temperature and centrifuged at 17,000 × g for 20 minutes to pellet residual Ti_3_C_2_T*_x_* MXenes. Supernatants containing the tryptic peptides were carefully transferred into fresh low-binding Eppendorf tubes. Formic acid (FA) was added to a final concentration of 5% (v/v) to acidify the samples and adjust pH to between 2 and 3, ensuring peptide stability and compatibility with desalting. Peptide desalting and cleanup were performed using C18 StageTips. Each tip was first activated with 80% acetonitrile (ACN) and 5% FA, equilibrated with 5% FA, and then loaded with the peptide-containing samples. Bound peptides were washed with 5% FA and eluted in two stages: first with 30% ACN + 5% FA, and then with 80% ACN + 5% FA. Both eluates for each sample were pooled, gently mixed, and transferred into a new low-binding tube. Samples were dried using a vacuum centrifuge (SpeedVac) and stored at –20 °C prior to shipment. Dried peptides were shipped overnight on dry ice to the core proteomics facility using FEDEX with next-morning delivery. Sample integrity upon arrival was confirmed by the receiving facility.

### Physicochemical characterizations

X-ray diffraction (XRD) patterns of the as-synthesized Ti_3_AlC_2_ MAX phase and Ti_3_C_2_T*_x_* MXenes were obtained using a XRDynamic 500 diffractometer from Anton Paar using Cu Kα (λ = 1.5406 Å). The samples for XRD measurements were mounted on the fixed sample stage and scanned from 5° to 70° with a step size of 0.01° and a time per step of 20 s. Raman spectrum of the Ti_3_C_2_T*_x_* MXene film was collected using XploRA PLUS Raman microscope from Horiba Scientific. For Raman spectra, 785 nm laser was used with a grating size of 1200 gr/mm and a power 0.1%.

SEM was performed using a JEOL JSM-7800f FESEM with a lower electron detector at an acceleration voltage of 5 kV and 15 kV to study the surface morphology and flake size. The Ti_3_C_2_T*_x_* MXene solution concentration was maintained at < 0.1 mg/mL and loaded on an anodic disc, followed by vacuum drying for 2 hours. The samples were gold sputtered to reduce the charging and improve the sharpness of the SEM images.

TEM was carried out using a JEM-120i (JEOL Ltd.) operated at 120kV. 20 μL of the bare Ti_3_C_2_T*_x_* MXene was deposited onto a copper grid and used for imaging. For PC-coated Ti_3_C_2_T*_x_* MXene, 20 μL of sample was negatively stained using 20 μl uranyl acetate 1%, washed with DI water, deposited onto a copper grid, and used for imaging. DLS and zeta potential analyses were performed to measure the size distribution and surface charge of the Ti_3_C_2_T*_x_* MXenes before and after PC formation using a Zetasizer Lab series DLS instrument (Malvern company). A Helium Neon laser with a wavelength of 632 nm was used for size distribution measurement at room temperature.

### LC-MS/MS analysis and proteomics data processing

Each dried vial was reconstituted with 22 µl 2% ACN 0.1% FA. 5 µL of each digest was injected and run 3 times by nanoLC-MS/MS using a 2h gradient on a Waters CSH 0.075mm x 250mm C18 column feeding into a Thermo Eclipse Tribrid Orbitrap mass spectrometer run in FT-IT mode, using typical settings with HCD for fragmentation. Raw mass spectrometry .raw files were converted to the mzML format using ThermoRawFileParser v.1.4.3. Database searches were performed using Sage v.0.14.7 against the human UniProt Swiss-Prot database downloaded from UniProt in September 2025.^60^ Precursor and fragment ion tolerance were set to ±20 ppm with the following modifications: carbamidomethylation of cysteine (+57.0215 Da, static) and oxidation of methionine (+15.9949 Da, variable). Peptides and proteins were filtered at a 1% FDR threshold.

Isoform collapsing and protein parsimony were performed using a custom R-based workflow: peptides mapping to canonical and non-canonical isoforms were collapsed to the canonical protein if not uniquely identifying an isoform, and shared peptides were removed when a non-canonical isoform had a unique peptide. Peptide-level intensities were aggregated to protein-level by summing across peptides, followed by variance stabilization normalization (VSN) using the limma package.^61^

Principal component analysis (PCA) was performed in RStudio using variance-stabilized protein intensity data generated through the limma-based workflow. Samples with low total signal or designated as technical controls were excluded from downstream analysis. Differential expression was conducted using limma v.3.64.3 in R v.4.5.1 with a design matrix with AD and healthy samples and subject sample as factors.^62^ Significance was determined using a limma FDR-adjusted p-value cutoff ≤ 0.05 and absolute log_2_FC ≥ 1. Additional analysis was performed with the aid of the ggplot, clusterProfiler, and pheatmap packages in R.

## Results and discussion

Prior to PC formation, the as-synthesized Ti_3_C_2_T*_x_* MXene nanosheets were characterized to evaluate their physicochemical properties. The physicochemical characterization of uncoated (bare) MXenes was conducted using XRD, Raman spectroscopy, DLS, zeta potential measurements, SEM, and TEM. **Figure 2A** shows the XRD pattern of as-synthesized Ti_3_AlC_2_ MAX and Ti_3_C_2_T*_x_* MXene nanosheets. Comparing the XRD patterns of Ti_3_AlC_2_ MAX (MXene precursor) with the Ti_3_C_2_T*_x_* MXene nanosheets shows the complete selective etching of the precursor and formation of the MXene sheets with only distinctive (00L) peaks and the first peak at ∼8°.^58^ Raman spectrum of the synthesized Ti_3_C_2_T*_x_* MXene nanosheets in **Figure 2B** reveals three characteristic regions (i) flake region, which corresponds to a collective vibration of carbon, the titanium layers, and surface groups, (ii) T*_x_* region corresponding to the vibrations of surface groups (-O, -OH, -F, -Cl) and (iii) carbon region, which encompasses both in-plane and out-of-plane vibrations of carbon atoms.^63^ Additionally, the absence of free carbon (D and G peak) indicates that the prepared Ti_3_C_2_T*_x_* MXene nanosheets are free of carbon impurities, which indicate high-quality MXene as the presence of free carbon could be due to the low-quality synthesis or MXene degradation due to hydrolysis-assisted oxidation.^63-64^

**Figure 2:**
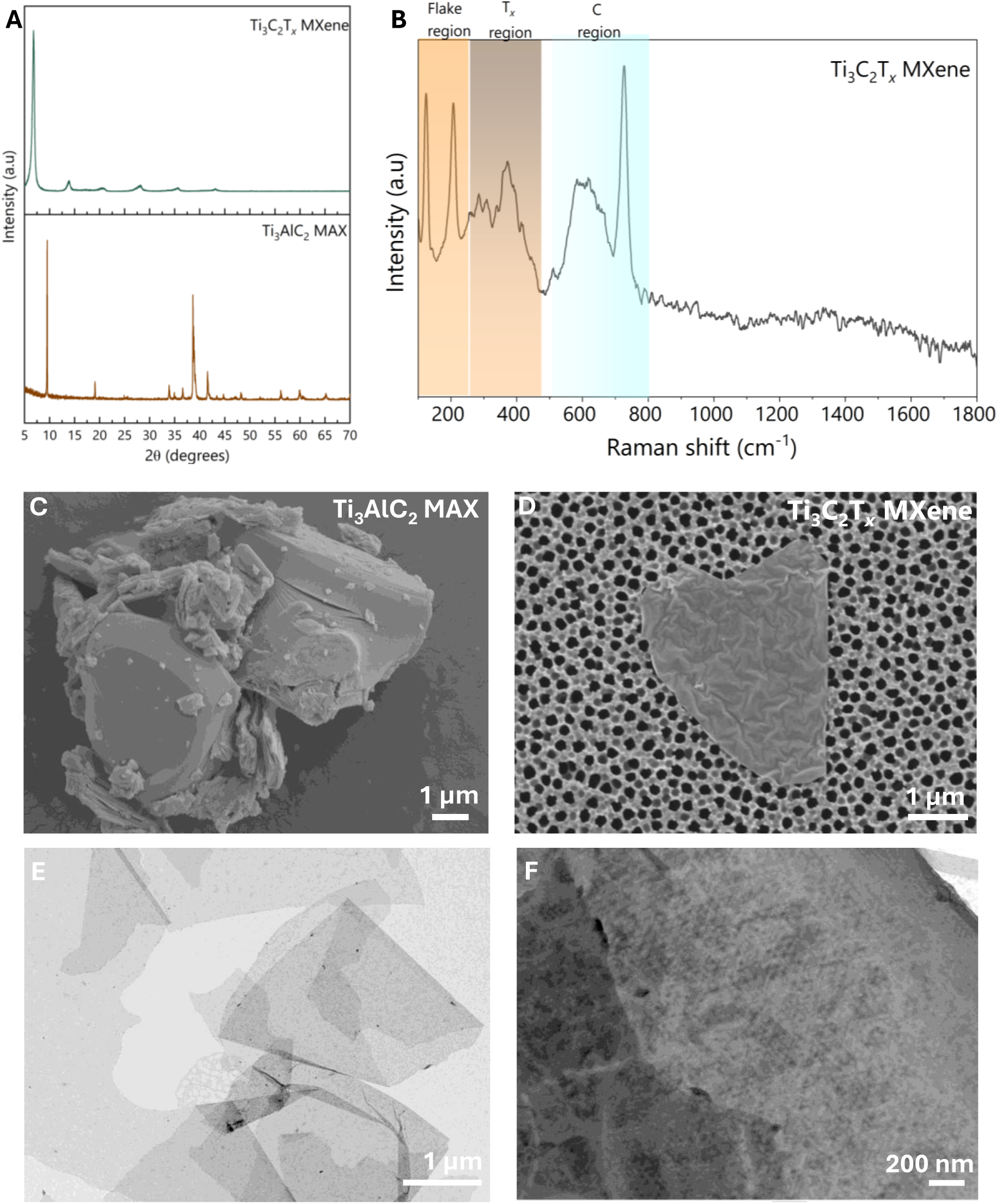
(A) XRD patterns of the MXene precursor (Ti_3_AlC_2_ MAX) and the as-synthesized Ti_3_C_2_T*_x_* MXene nanosheets, (B), Raman spectrum of Ti_3_C_2_T*_x_* MXene with labeled different regions, SEM images of (C) Ti_3_AlC_2_ MAX and (D) Ti_3_C_2_T*_x_* MXene nanosheets, (E) TEM image of bare MXenes, and (F) TEM images of PC-coated MXenes following incubation with a healthy control plasma sample, as representative.

Next, the synthesized Ti_3_AlC_2_ MAX (MXene precursor) and Ti_3_C_2_T*_x_* MXene samples were characterized using microscopic techniques. SEM micrograph of the Ti_3_AlC_2_ MAX shown in **Figures 2C** and **2D** reveals the layered morphology of the bulk carbide structures. The SEM micrograph of Ti_3_C_2_T*_x_* MXene in **Figure 2D** represents a single-to-few-layered Ti_3_C_2_T*_x_* MXene flake or nanosheet. The TEM image of as-synthesized MXenes (**Figure 2E**) clearly confirms the ultrathin, 2D morphology of the exfoliated nanosheets, exhibiting high transparency under the electron beam. The sheets show smooth surfaces and sharp edges, consistent with minimal oxidation or aggregation following synthesis. After incubation with human plasma (from a healthy control as representative), the TEM image of the Ti_3_C_2_T*_x_* MXene–PC complex reveals a thin, diffuse organic coating surrounding the nanosheet surfaces (**Figure 2F**). The PC layer is visible as a slightly darker or rougher contrast zone on the MXenes flakes, indicating successful adsorption of plasma proteins. ^65-67^ In fact, the presence of this coating confirms the formation of a stable hard PC, which alters the surface chemistry, hydrodynamic behavior, and colloidal stability of Ti_3_C_2_T*_x_* MXenes in biological environments. These structural features provide a large accessible surface area and active surface terminations (–O, –OH, –F) conducive to protein adsorption and charge transport, which are essential for biosensing applications.^68-70^

DLS analysis revealed that the as-synthesized bare Ti_3_C_2_T*_x_* MXenes have an average hydrodynamic diameter of ∼ 1100 nm and a surface zeta potential of ∼ –36 mV (**Figure 3** and **Table 1**). Upon incubation with human plasmas, the Ti_3_C_2_T*_x_* MXenes underwent noticeable surface remodeling: the average size increased to about 1120-1275 nm, and the surface charge shifted to –27 to –20 mV approximately, reflecting the successful formation of a PC layer (see **Figure S1** and **Table S1** of SI). Since DLS provides the hydrodynamic diameter rather than the true lateral flake dimensions of two 2D MXene nanosheets, the estimated MXene flake size was calculated using the previously reported empirical relationship developed for Ti_3_C_2_T*_x_* MXenes systems:^58^

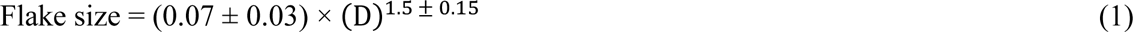

**Figure 3:**
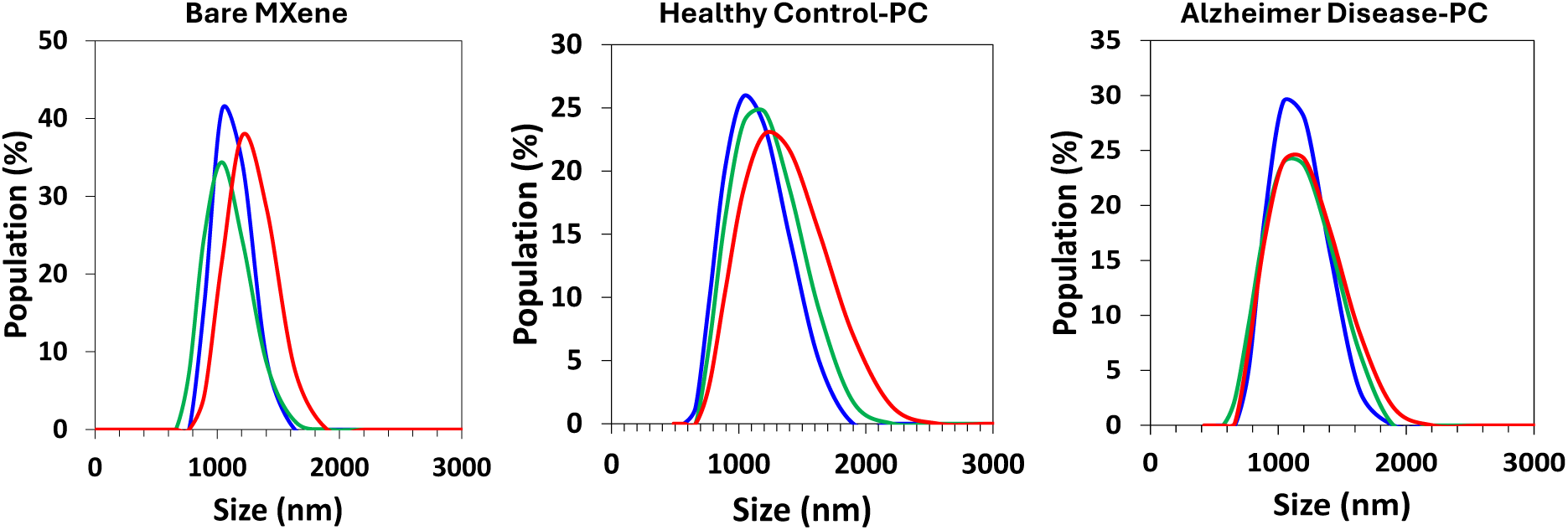
Representative DLS analysis of Ti_3_C_2_T_*x*_ MXene–PC complexes incubated with plasma from Alzheimer’s disease patients and healthy controls (see SI for the full analysis of the healthy and disease samples). Each curve represents the hydrodynamic size distribution of 2D Ti_3_C_2_T_*x*_ MXene nanostructures following incubation with individual plasma samples. Different colors correspond to three independent repeats.

**Table 1:** Summary of average size, polydispersity index (PDI), zeta potential, and the corresponding standard deviation (SD) values of MXene–PC complexes formed with plasma from Alzheimer’s disease patients and healthy controls, as representative.

| MXene–PC Sample | DLS hydrodynamic Size (nm) | SD (nm) | PDI | DLS flake Size (nm) | SD (nm) | SD | Zeta potential (mV) | SD (mV) |
| --- | --- | --- | --- | --- | --- | --- | --- | --- |
| Bare MXene | 1095.7 | 80.1 | 0.584 | 2539 | 278 | 0.07 | -35.7 | 0.6 |
| Healthy Control-PC | 1128.1 | 114.3 | 0.407 | 2653 | 403 | 0.075 | -25.0 | 0.7 |
| Alzheimer Disease-PC | 1152.3 | 80.1 | 0.372 | 2740 | 276 | 0.037 | -21.6 | 0.1 |

Where D is the hydrodynamic size of the flakes obtained by DLS. Based on equation 1, the estimated flake sizes were approximately 2.54 ± 0.28 µm for bare MXene, 2.65 ± 0.40 µm for HC PC, and 2.74 ± 0.29 µm for AD PC samples. Two different Alzheimer’s disease and HC groups are denoted by AD and HC, and numbers 1 to 10 in **Figure S1** and **Table S1** refer to 10 different individual samples per each disease and healthy group. Next, plasma samples from AD patients and healthy individuals were collected and incubated with Ti_3_C_2_T*_x_* MXene nanosheets to form MXene–PC complexes. The average hydrodynamic diameter, polydispersity index (PDI), and zeta potential values of all samples are summarized in **Table S1**. The formation of the PC on Ti_3_C_2_T*_x_* MXene surfaces resulted in a noticeable increase in hydrodynamic size and PDI, along with a decrease in surface charge, reflecting the successful adsorption of plasma proteins and the consequent alteration of surface behavior.

Figure 4 presents SDS-PAGE profiles of proteins adsorbed onto Ti_3_C_2_T*_x_* MXene nanosheets following incubation with plasma from healthy controls (HC1–HC10) and Alzheimer’s disease patients (AD1–AD10). Distinct banding patterns were observed between the two groups, indicating differences in the molecular composition of the Ti_3_C_2_T*_x_* MXene-bound PC. Across all samples, proteins spanning a broad molecular weight range (∼10–250 kDa) were detected, consistent with adsorption of abundant plasma constituents such as albumin (∼66 kDa), immunoglobulins (∼150 kDa), apolipoproteins (20–40 kDa), and complement-associated proteins. Notably, AD-derived PC samples exhibited stronger staining intensity and broader signal distribution in the ∼50–100 kDa range, suggesting enhanced association of mid-to-high molecular weight proteins.

**Figure 4:**
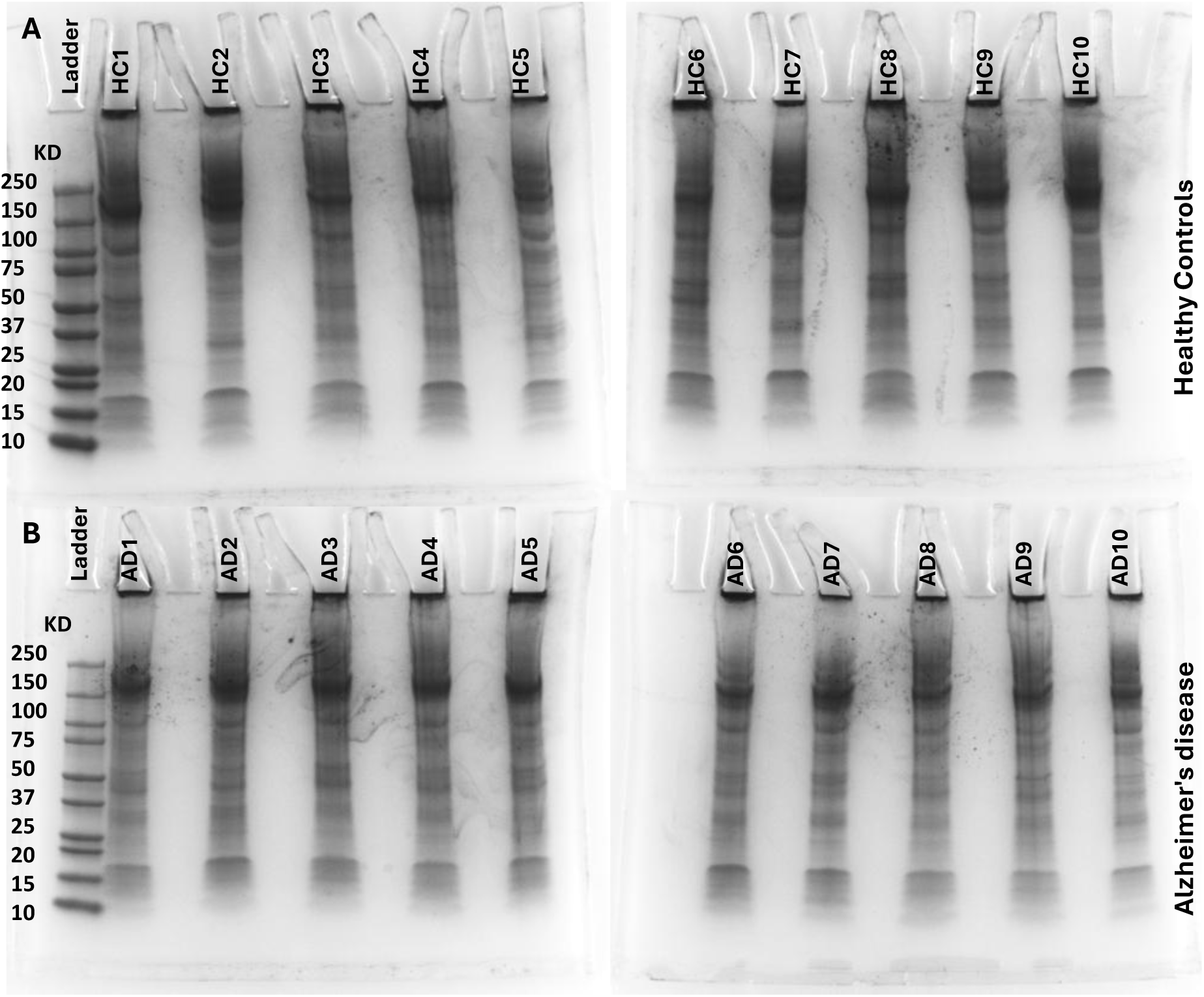
SDS-PAGE analysis of PC compositions formed on Ti_3_C_2_T_*x*_ MXene nanosheets after incubation with plasma from AD patients and HC. Lanes correspond to individual plasma samples (n = 10 per group), showing distinct banding patterns that reflect variations in protein adsorption profiles. Differences in band intensity and distribution indicate group-specific PC compositions between (a) HC-and (b) AD- derived Ti_3_C_2_T_*x*_ MXene–PC complexes.

Several AD lanes also displayed increased smearing toward higher molecular weights, indicative of a more heterogeneous corona composition. In contrast, HC-derived samples showed more defined and reproducible banding patterns, with relatively higher intensity in the lower molecular weight region (e.g., 20–40 kDa), consistent with a more homogeneous protein adsorption profile. These qualitative differences support the presence of disease-dependent Ti_3_C_2_T*_x_* MXene–PC fingerprints, reflecting altered plasma protein composition and protein–surface interactions in AD. Additionally, these qualitative observations are consistent with the downstream label-free proteomics results (see next section), which revealed statistically significant differential enrichment of a subset of proteins in AD-derived Ti_3_C_2_T*_x_* MXene coronas relative to HC.

Unlike direct plasma proteomics, which is dominated by a small number of highly abundant proteins, PC–based proteomics reflects a selectively enriched sub-proteome shaped by competitive adsorption at the nano–bio interface.^71-72^ The Ti_3_C_2_T*_x_* MXene surface acts as a physicochemical filter, amplifying proteins with higher surface affinity, specific charge distributions, and structural motifs.^73-74^ As a result, the resulting PC proteome (Figure 5) provides a compressed and information-rich molecular representation of plasma alterations associated with AD. ^75^ In order to find group-specific protein enrichment patterns, hierarchical clustering was performed on normalized label free quantification (LFQ) intensities of the differentially expressed proteins (Figure 5A), which shows partial segregation between AD-and HC-derived Ti_3_C_2_T*_x_* MXene–PC along the x-axis (sample axis), where columns corresponding to AD samples cluster separately from HC. This separation is reflected in distinct color patterns (z-scored LFQ intensities), where groups of proteins exhibit consistently higher (blue) or lower (yellow/red) expression across one condition relative to the other. Notably, clusters of proteins on the y-axis (protein axis) show coordinated expression changes that differentiate AD from healthy samples, supporting disease-associated proteomic signatures.

**Figure 5:**
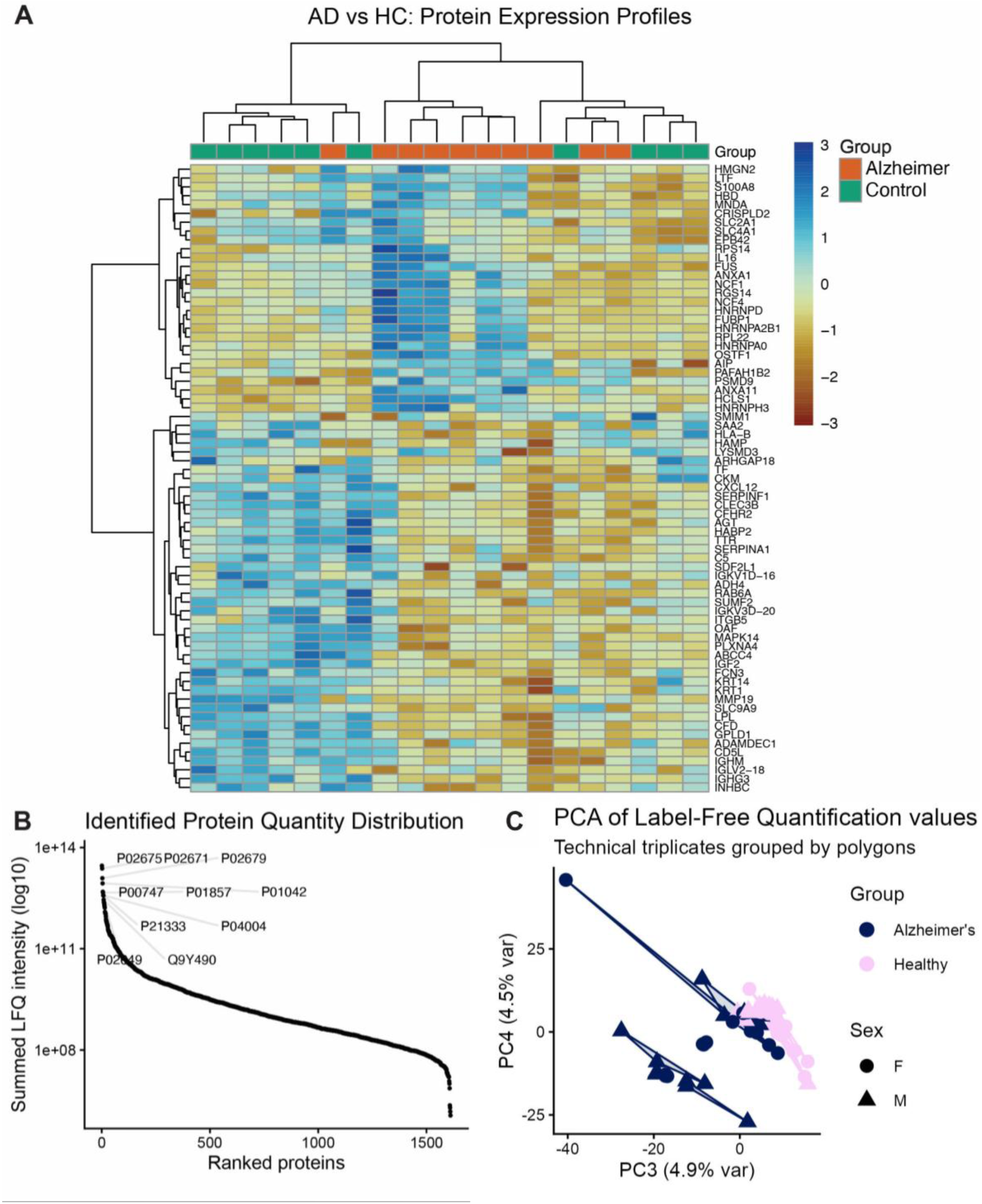
(A) Heatmap of differentially expressed proteins identified in Ti_3_C_2_T_*x*_ MXene–PC following incubation with AD and healthy plasma samples. (B) A total of 1611 proteins were quantified across samples. (C) PCA clustering of Ti_3_C_2_T*_x_* MXene–PC samples grouped by sex and disease condition. Technical triplicates are denoted by lines connecting them.

The heatmap in Figure 5A also demonstrates that inter-individual variability within each group is substantially lower than between-group variation, indicating the robustness of the MXene–PC interface in capturing disease-associated proteomic signatures. The distinct clustering pattern suggests that Ti_3_C_2_T*_x_* MXene does not merely adsorb abundant plasma proteins indiscriminately but instead enriches distinct molecular subsets reflective of the underlying pathological state. We performed a comprehensive proteomics analysis of the PC profiles for each group, identifying a total of 1,611 proteins (Figure 5B and **Figure S2**). Figure 5B demonstrates the broad dynamic range of proteins identified within the MXene–PC samples, including both high- and low-abundance plasma proteins. This is important because many biomarkers are typically present at low concentrations in biological samples. Importantly, several identified proteins are known AD-associated markers or potential novel biomarker candidates, indicating the ability of the MXene platform not only for disease discrimination but also for biomarker discovery and molecular fingerprinting applications.

To quantitatively compare the PC proteome profiles, the mass spectrometry intensities of proteins were normalized using VSN with the limma R package to improve comparability across various LC-MS runs. ^62^ PCA of the variance-stabilized protein data shows clustering by individual samples (Figure 5C). Because PC1 and PC2 were partially associated with demographic variables, including age, additional PCA analysis focusing on PC3 and PC4 was performed to better visualize disease-related proteomic differences independent of demographic effects. Additional PCA demographic association analysis is provided in **Figure S3 o**f the SI. Each point represents an individual sample (with technical replicates connected), and the spread of points within each group reflects inter-individual variability in PC composition. AD-derived samples exhibit a broader dispersion compared to HC, indicating greater heterogeneity, likely due to the complexity of AD and variation in patient demographics. Despite this variability, PCA reveals a partial but reproducible separation between AD- and HC-derived Ti_3_C_2_T*_x_* MXene–PC proteomes along the principal components, supporting disease-associated differences. These findings suggest that disease-associated proteomic variance is captured more prominently in PC3 and PC4 after accounting for demographic-related variability present in the earlier principal components.

Importantly, the persistence of disease-associated clustering across technical replicates highlights the robustness of the Ti_3_C_2_T*_x_* MXene–PC interface in capturing shared molecular features of AD, even in the presence of biological variability.^76-77^ Application of stringent statistical filtering criteria (adjusted P < 0.05 and log_2_ fold change>1), identified 71 significantly differentially enriched proteins between AD and HC studies (**Figure S2**). Although this represents a selective subset of the quantified proteome, these proteins form a coherent disease-associated signature rather than isolated changes. The relatively modest number reflects the use of conservative statistical thresholds and highlights that only the most robust alterations in plasma composition are transduced at the Ti_3_C_2_T*_x_* MXene–PC interface. Importantly, the detection of a focused panel of statistically significant proteins suggests that Ti_3_C_2_T*_x_* MXene–PC profiling enhances signal specificity while minimizing stochastic adsorption effects, thereby enabling discrimination between AD and healthy samples.

Conventional single NPs’ PC workflows typically report on the order of hundreds to several hundred quantified proteins, although values vary substantially with particle chemistry, sample pooling strategy, and proteomic workflow.^78-82^ For example, PC on conventional organic and inorganic (e.g., silica, polystyrene, and gold) NPs have been reported to contain more than 250 proteins, while more recent single-NP workflows reported average quantified depths of ∼600 for polystyrene and ∼900 for modulated PC of polystyrene NPs.^78, 83-85^ Beyond conventional spherical NPs, several 2D nanomaterials have also been shown to form distinct and biologically relevant PC.^74, 86-88^ Studies on graphene demonstrate that it can adsorb hundreds of plasma proteins (typically ∼300–400 depending on structure and conditions), with PC composition strongly influenced by surface chemistry, layer number, and biological environment.^87, 89-90^ Importantly, graphene-based systems have been used to generate personalized, disease-dependent PC profiles from human plasma, highlighting their potential for diagnostic applications.^87^ Similarly, MXene nanosheets have been shown to interact with human plasma proteins, exhibiting selective enrichment of immunoglobulins, coagulation factors, and complement-related proteins, indicating a strong and selective nano–biointerface.^74^

Other 2D materials such as borophene and MoS_2,_ also form PC, although typically with lower reported proteome depth (∼90–100 proteins) and with a stronger emphasis on immune and surface interaction effects.^88, 91^ Overall, these studies suggest that while 2D nanomaterials consistently exhibit high surface reactivity and selective protein adsorption, their use in deep, large-scale plasma proteome profiling remains relatively limited, indicating the potential of MXene-based platforms for enhanced PC–enabled biomarker discovery.^92^ It is noteworthy that Ti_3_C_2_T*_x_* MXene–PC in the present study enabled the detection of over 1600 proteins across the cohort, demonstrating substantially enhanced proteomic coverage (**Figure S2**). This increased depth likely arises from the synergistic combination of high surface-to-volume ratio, 2D planar geometry, and abundant surface terminations (–O, –OH, –F), which collectively promote extensive and diverse protein adsorption.

Differential protein expression between AD and HC is shown in Figure 6A. Fold changes were calculated as log₂(AD/HC), such that positive values indicate enrichment in AD and negative values indicate enrichment in HC. The x-axis represents log_2_ fold change (where ±1 corresponds to a twofold difference), and the y-axis represents log_10_ (adjusted *p*-value). Applying thresholds of adjusted *p* < 0.05 and |log_2_FC| > 1, a subset of proteins was identified as significantly differentially enriched between the two groups. Several heterogeneous nuclear ribonucleoproteins (hnRNPs) were identified as upregulated in the AD population compared to the control (hnRPA2B1, hnRNPD, hnRNPH, hnRNPA0), as depicted in Figure 6A. HnRNPs are ubiquitously expressed proteins that play a role in various cellular processes, including pre-mRNA processing, RNA shuttling to the cytoplasm, and miRNA sorting. ^93^ We found that HNRNPA2B1 protein abundance was ∼2.4-fold higher in the AD samples (Figure 6A); this is consistent with previous reports showing that HNRNPA2B1 links oligomerized tau protein to N⁶-methyladenosine–modified RNA transcripts.^94^ Notably prior studies have reported that the formation of this complex can increase by up to ∼5-fold in human AD subjects.^94^ Circulating HNRNPA2B1 has also been previously linked to cognitive impairment, found in higher concentrations in patients suffering from postoperative neurocognitive dysfunction.^95^ The enrichment of hnRNP family members suggests that Ti_3_C_2_T*_x_* MXene surfaces preferentially capture proteins involved in RNA processing and stress granule dynamics, processes increasingly implicated in tau pathology and neurodegeneration, and therefore warrants further investigation.^96-97^

**Figure 6:**
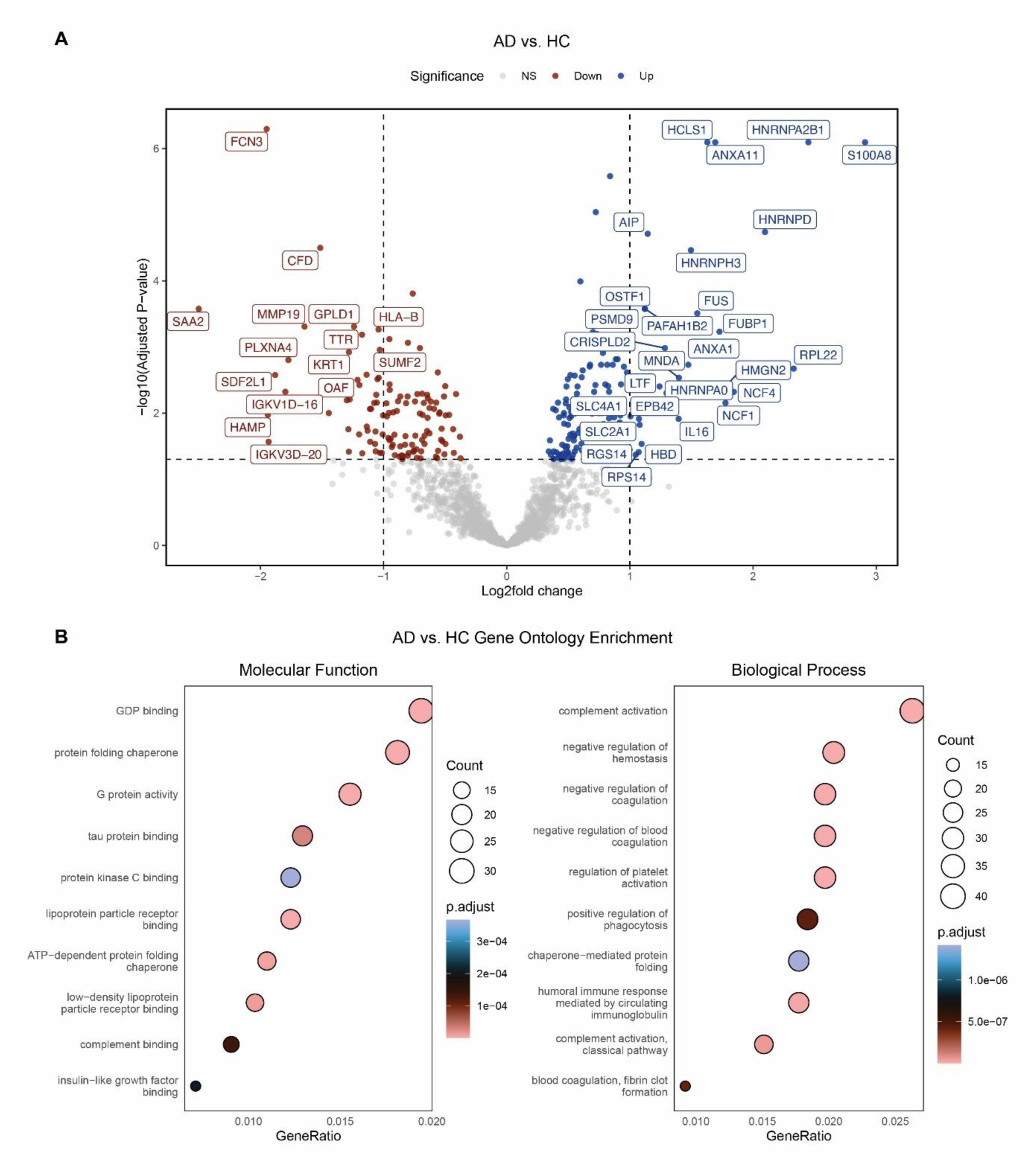
(A) Differentially regulated proteins purified from patient PC’s, labeled proteins met a log_2_ fold change minimum of 1 and adj. P value of < 0.05 to be labeled “significant”. (B) gene ontology enrichment analysis results for molecular function and biological processes overrepresented by significantly upregulated proteins.

Moreover, calgranulin-A, or protein S100A8, was found significantly upregulated in AD subjects (2.9-fold) over HC subjects (Figure 6A). An inflammatory mediator and calcium-binding protein, previous reports have demonstrated S1008A’s expression is significantly increased in the hypothalamic of two AD mouse models.^98-99^ Previous studies have reported inconsistent changes in circulating S100A8 levels in AD, indicating that further investigation is required to clarify its potential as a plasma biomarker. The identification of inflammatory mediators and complement-associated proteins aligns with extensive evidence linking systemic immune activation and complement dysregulation to AD progression, supporting the biological relevance of the Ti_3_C_2_T*_x_* MXene–PC proteome. Additionally, annexin proteins A1 and A11 (ANXA1, ANXA11) were both upregulated in AD serum, ∼1.7 and 1.47-fold, respectively (Figure 6A). Annexin proteins have well-described roles in Ca^2+^-regulated phospholipid binding and membrane trafficking, as well as more recently uncovered roles in sensing cellular stress via extracellular functions.^100^

Another protein identified in our differential analysis (Figure 6A), ANXA1 has been implicated in regulating blood-brain-barrier (BBB) integrity, and recombinant ANXA1 treatment has been demonstrated to have protective effects against Aβ42-induced BBB disruption.^101^ Notably, previous reports have found decreased levels of ANXA1 in AD patient sera, contrary to our findings.^101^ Another annexin family member identified in our differential analysis (Figure 6A), ANXA11, has been reported to interact with TDP-43, a key pathological protein frequently mislocalized and aggregated in AD.^102^ Annexin proteins lack classical signal sequences directing them for secretion, thus, the regulation of their escape into circulation is poorly understood.^100^ Annexins are not classical plasma biomarkers, and their enrichment in the Ti_3_C_2_T*_x_* MXene–PC suggests that non-canonical circulating proteins, potentially released through vesicles or cell stress pathways, are preferentially captured by the nanointerface.^103-104^ The full protein list can be found in **Supplementary Table S3**.

To further interpret these differences, gene ontology (GO) enrichment analysis was performed on the significantly altered proteins (Figure 6B), revealing overrepresented biological processes and molecular functions associated with AD-related proteomic changes. The GO enrichment analysis of proteins upregulated in AD-derived PCs revealed significant enrichment in tau protein binding and protein folding and quality control pathways, both of which are critically disrupted in AD.^105^ Analysis of biological process GO terms further indicated a pronounced inflammatory signature, with complement activation emerging as a dominant pathway.^106^ The overrepresentation of tau-binding, proteostasis-related, and complement activation pathways suggests that Ti_3_C_2_T*_x_* MXene–PC profiling captures molecular processes key to protein misfolding, immune activation, and loss of proteome homeostasis in AD, rather than isolated biomarker changes.^12, 107^ These findings indicate that Ti_3_C_2_T*_x_* MXene nanosheets provide an enriched molecular interface capable of capturing a broader spectrum of plasma proteins compared to conventional nanoparticles, thereby expanding the analytical window for disease-associated biomarker discovery.

## Conclusions

In summary, this work demonstrates that Ti_3_C_2_T*_x_* MXene–PC complexes provide a robust nanointerface for capturing disease-associated plasma signatures in AD. Physicochemical measurements revealed that PC formation consistently altered the hydrodynamic behavior and surface charge of Ti_3_C_2_T*_x_* MXene in a way that reflected the underlying biofluid environment, demonstrating the sensitivity of this platform to plasma state. Importantly, proteomic profiling of the isolated PC layers enabled the quantification of 1,611 proteins without additional fractionation, highlighting the capacity of Ti_3_C_2_T*_x_* MXene–PC interfaces to enrich and reveal molecular features that are often masked in conventional plasma analyses. At the molecular level, AD-derived PC showed selective enrichment of protein classes linked to RNA regulation (including hnRNP family members), membrane stress and trafficking (annexins), and inflammatory signaling, pointing to coordinated biological processes consistent with AD pathophysiology rather than isolated single-analyte changes. Our findings suggest Ti_3_C_2_T*_x_* MXene–PC profiling as a versatile strategy for nanomaterial-assisted molecular phenotyping of neurodegenerative disease states, which provides an enriched molecular interface capable of capturing a broader spectrum of plasma proteins. Future studies will focus on expanding cohort size, associating PC signatures with clinical variables and disease stage, and translating these signatures into streamlined readouts through integration with complementary sensing modalities.

## Associated content

### Supporting Information

Supporting figures/tables and LC-MS/MS analyses.

### Notes

The authors declare no competing financial interest.

### Data Availability

The raw mass spectrometry data associated with this project have been uploaded to ProteomeXchange at PXD074560 via JPOST^108^ (Reviewer access https://repository.jpostdb.org/preview/1855276536994c537c8662 Access key 5582) .

## Acknowledgments

S.V and A.A.A acknowledge financial support from the National Institutes of Health (grant number R03EB034817). E.L. is supported in part by the University of Colorado School of Medicine Translational Research Scholar Program (TRSP) and the Research Investment in the Scientific Enterprise (RISE) awards. A.T. and B.A. acknowledge the support of the U.S. National Science Foundation, award number CMMI-2134607.

