## Supporting Information for "MXene–Protein Corona Interfaces for Molecular Profiling of Alzheimer’s Disease"

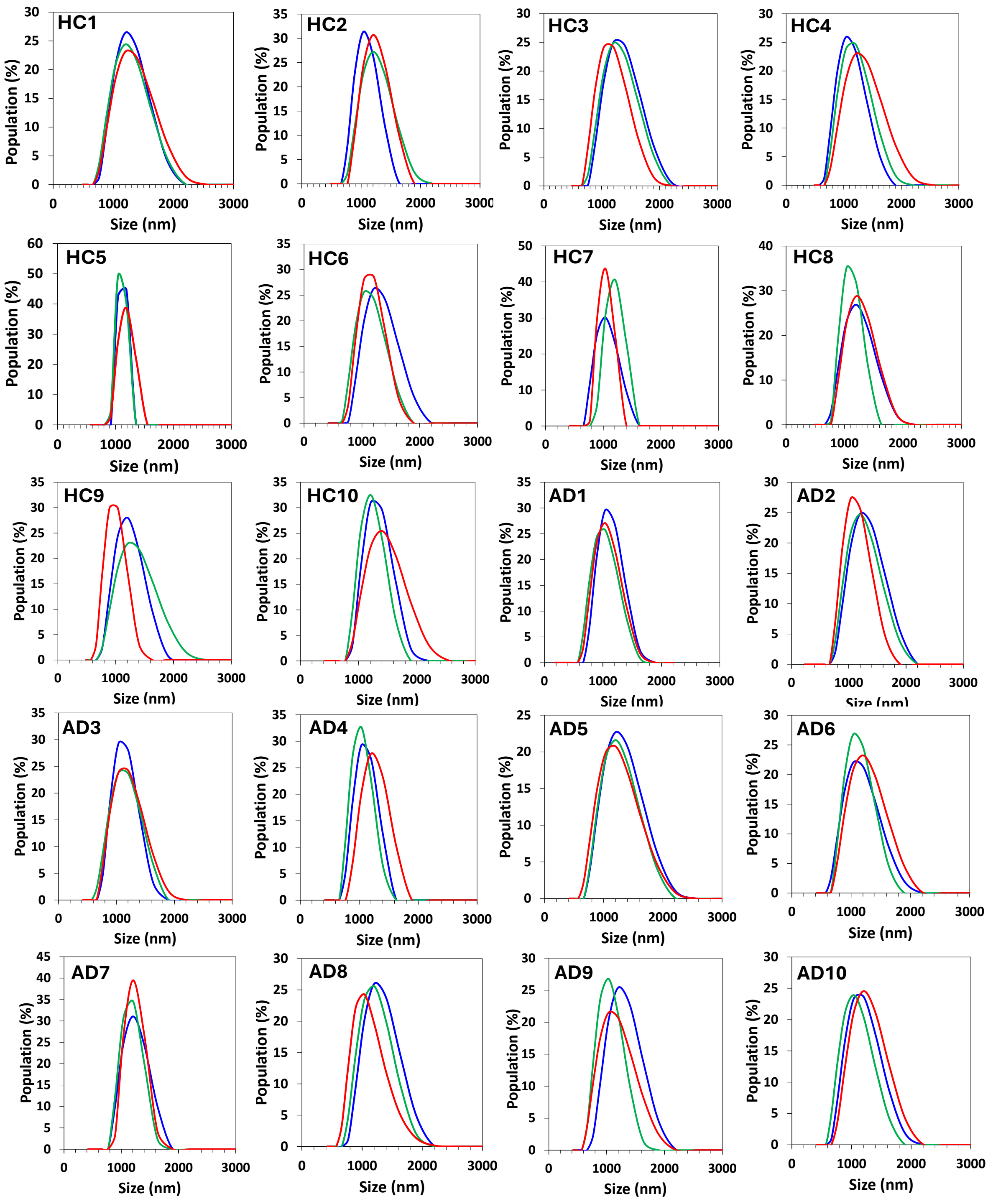


**Figure S1:** DLS analysis of $Ti_3C_2T_x$ MXene–PC complexes incubated with plasma from Alzheimer's disease (AD) patients and healthy controls (HC). Each curve represents the hydrodynamic size distribution of 2D $Ti_3C_2T_x$ MXene nanostructures following incubation with individual plasma samples ($n = 10$ per group). Different colors correspond to three independent repeats.

**Table S1:** Summary of average size, polydispersity index (PDI), zeta potential (n = 10 per group, three technical replicates per sample), and the corresponding standard deviation (SD) values of MXene–PC complexes formed with plasma from AD patients and HC.

| MXene–PC Sample | DLS hydrodynamic Size (nm) | SD (nm) | PDI | DLS flake Size (nm) | SD (nm) | SD | Zeta potential (mV) | SD (mV) |
|---|---|---|---|---|---|---|---|---|
| **Bare** | 1095.7 | 80.1 | 0.584 | 2538.8 | 278.4 | 0.07 | -35.7 | 0.6 |
| **HC-PC-1** | 1272.7 | 24.7 | 0.289 | 3178.2 | 92.5 | 0.063 | -24.2 | 0.3 |
| **HC-PC-2** | 1176.5 | 73.3 | 0.399 | 2824.8 | 264 | 0.014 | -21.6 | 1.2 |
| **HC-PC-3** | 1253.8 | 69.1 | 0.378 | 3107.7 | 256.9 | 0.137 | -27.6 | 0.1 |
| **HC-PC-4** | 1191.5 | 81.9 | 0.407 | 2879 | 296.8 | 0.075 | -25.0 | 0.7 |
| **HC-PC-5** | 1173.1 | 45.5 | 0.323 | 2812.6 | 163.6 | 0.016 | -24.0 | 0.4 |
| **HC-PC-6** | 1187.0 | 78.3 | 0.303 | 2862.7 | 283.3 | 0.037 | -24.4 | 0.7 |
| **HC-PC-7** | 1202.3 | 117.6 | 0.366 | 2918.2 | 428.2 | 0.038 | -24.6 | 0.1 |
| **HC-PC-8** | 1192.8 | 62.8 | 0.331 | 2883.7 | 227.7 | 0.058 | -22.0 | 1.1 |
| **HC-PC-9** | 1164.3 | 139.3 | 0.399 | 2781 | 499.1 | 0.041 | -24.0 | 0.1 |
| **HC-PC-10** | 1274.6 | 83.9 | 0.688 | 3185.4 | 314.5 | 0.011 | -19.4 | 0.6 |
| **AD-PC-1** | 1152.5 | 38.5 | 0.407 | 2738.8 | 137.2 | 0.065 | -23.7 | 0.2 |
| **AD-PC-2** | 1206.9 | 67.1 | 0.418 | 2935 | 244.8 | 0.042 | -23.9 | 0.6 |
| **AD-PC-3** | 1125.4 | 21.1 | 0.372 | 2642.8 | 74.3 | 0.037 | -21.6 | 0.1 |
| **AD-PC-4** | 1121.5 | 89.8 | 0.399 | 2629 | 315.8 | 0.015 | -25.5 | 0.2 |
| **AD-PC-5** | 1235.4 | 31.2 | 0.314 | 3039.6 | 115.1 | 0.026 | -20.6 | 0.1 |
| **AD-PC-6** | 1156.6 | 51.1 | 0.473 | 2753.4 | 182.5 | 0.050 | -26.3 | 0.3 |
| **AD-PC-7** | 1204.3 | 27.4 | 0.380 | 2925.5 | 99.8 | 0.048 | -23.4 | 0.5 |
| **AD-PC-8** | 1186.1 | 83.4 | 0.494 | 2859.4 | 301.6 | 0.102 | -25.7 | 0.5 |
| **AD-PC-9** | 1143.2 | 93.4 | 0.691 | 2705.7 | 331.6 | 0.037 | -22.1 | 0.1 |
| **AD-PC-10** | 1152.7 | 74.4 | 0.430 | 2739.5 | 265.2 | 0.116 | -23.00 | 0.5 |

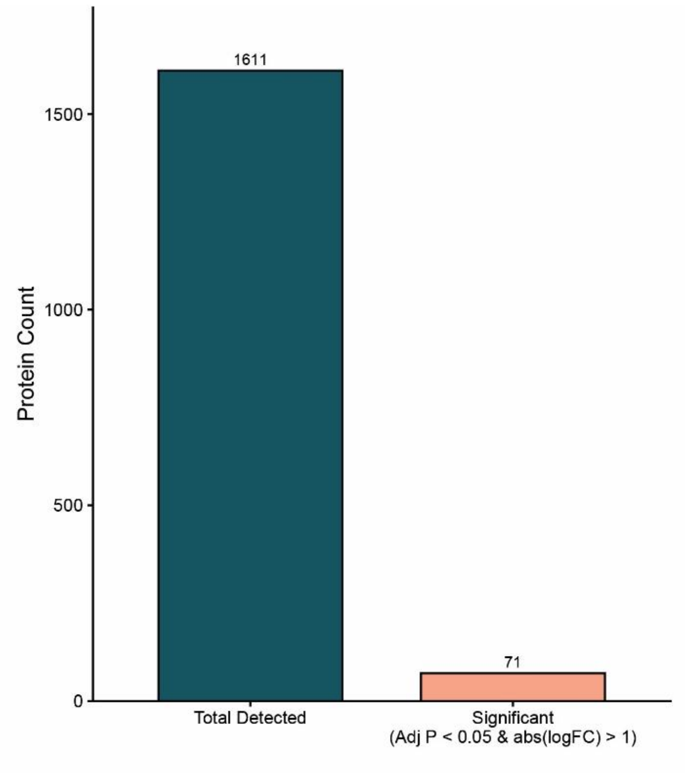


**Figure S2:** Summary of proteomic quantification results, including total proteins detected across samples and statistically significant different proteins (adjusted $P < 0.05$ and $\log_2$ fold change>1).

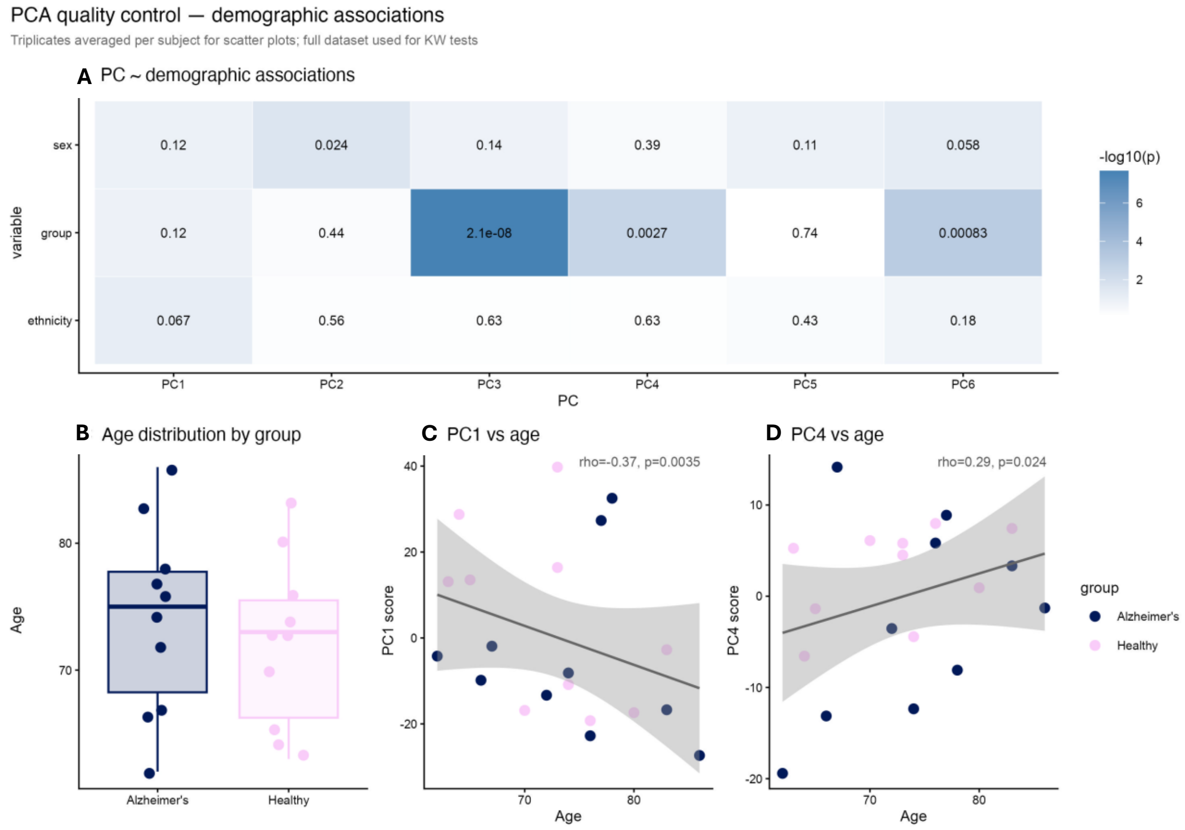


**Figure S3:** PCA demographic association analysis of MXene–PC proteomics data. (A) Heatmap showing statistical associations between principal components (PC1–PC6) and demographic variables including disease group, sex, and ethnicity. PC3 and PC4 showed the strongest association with disease group. (B) PCA quality control and demographic distribution analyses including age distribution and correlation of principal component scores with age.
